# Cultural transmission and biological markets

**DOI:** 10.1101/083907

**Authors:** Claude Loverdo, Hugo Viciana

## Abstract

Active cultural transmission of fitness-enhancing behavior can be seen as a costly strategy, one whose evolutionary stability poses a Darwinian puzzle. In this article, we offer a biological market model of cultural transmission that substitutes or complements existing kin-selection based theories for the evolution of cultural capacities. We explicitly formulate how a biological market can account for the evolution of deference and prestige-related phenomena, as well as how it can affect the dynamics of cumulative culture. We show that, under certain conditions, teaching evolves even when innovations are not sufficiently opaque and can be acquired by emulators via inadvertent transmission. Furthermore, teaching in a biological market is a precondition for enhanced individual learning abilities.

## 1. Introduction

A certain view of cultural evolution sees cultural transmission as bringing a straightforward advantage to the group or the individual’s kin. Hence, its evolution follows. However, if the active transmission of culture is such a successful strategy, then where is all the evidence of it in the animal kingdom? Although social learning and certain forms of animal traditions are common in many non-human species, active cultural transmission or teaching is a far rarer phenomenon (Boyd and Richerson, 1996; Thornton and Raihani, 2008).

On the production side of active cultural transmission, “natural pedagogy” —the dispositions and efforts of adults to make themselves easily understood by children and to thus facilitate the transmission of cultural knowledge— is certainly part of the human pattern of cultural transmission (Hewlett et al., 2011), a good candidate for a universal trait of our species, and perhaps even a biological adaptation (Csibra and Gergely, 2009). Such considerations suggest a vertical-transmission view of the evolution of human culture, i.e: direct transmission from the parental generation to the generation of siblings. Nonetheless, another view widely accepted among ethnographers claims that adult-infant instruction is rare in hunter-gatherers groups (Atran and Sperber, 1991). Moreover, as several case studies in cultural transmission have indicated, non-vertical transmission —that is, transmission to children from other children or slightly older individuals, as opposed to much older adults— is far more important for cultural transmission than assumed (Aunger, 2000; Morin, 2015). It has even been argued that non-vertical transmission might constitute the key component in children’s and young adults’ adoption of much of the cultural repertoire (Harris, 1998).

From a population genetics perspective, other considerations also counter the all-importance of vertical transmission in the evolution of culture. Cultural capabilities were plausibly “built for speed” and adaptability (Richerson and Boyd, 2000). However, pure vertical cultural transmission is more analogous to genetic adaptation, and thus has fewer of those properties: it is more often subject to maladaptive lag and inertia than other forms of cultural transmission (McElreath and Boyd, 2008). In changing environments, *mother does not always know best.* The facilitation of cultural transmission via genetic relatedness, namely as a form of evolved nepotism, is possibly part of the picture yet can be easily exaggerated. There are conflicts of interests between parents and siblings (Trivers, 1974). In principle, parental manipulation could be selected for, which in return could prompt the evolution of devices that counteract the effects of vertical cultural transmission among siblings (Trivers, 2011).

Active cultural transmission is essentially problematic on the grounds of its cost-benefit structure. If what an individual learns is so useful in terms of fitness that acquiring it makes sense for other individuals, then why bother actively transmitting it? From the standard inclusive fitness perspective of evolution, it follows that traits that do not benefit kin need to benefit their carriers in order to evolve by way of natural selection (Dessalles, 2001, 2006). However, a great deal of cultural transmission is both not directed to kin and costly enough to pose a Darwinian puzzle. The question thus remains: Why transmit culturally?

Seemingly altruist cultural transmission of fitness-enhancing information yields a free rider problem structured similarly to the standard prisoner’s dilemma. Briefly, since active cultural transmission of fitness-enhancing information (“teaching”) is a form of cooperation, every individual would be better off if other individuals cooperate, while he or she does not cooperate. Therefore, all else being equal, a population of individuals capable of cultural transmission could be expected to evolve toward a sub-optimal equilibrium, one in which cultural transmission is simply not practiced.

The ethological definition of teaching characterized it as a form of altruism from early on (Caro and Hauser, 1992). In principle, ecological conditions linked to kin selection and alloparentality might have facilitated the evolution of certain cultural capacities (Hrdy, 2009; Flinn and Ward, 2005). Thus, the immense majority of formal models which have been used to investigate the evolution of teaching have relied on genetic relatedness to explain its stability (Castro and Toro, 2014; Fogarty et al., 2011). Nevertheless, the abovementioned theoretical and empirical considerations largely justify the exploration of complementary, if not alternative, evolutionary pathways by which cultural capacities can reach an adaptive equilibrium in a given population.

In this article, we analyze conditions of the evolution of cultural transmission capacities in a biological market model. Originally proposed by behavioral ecologists Ronald Noë and Peter Hammerstein, biological markets arise when associations between biological individuals are sufficiently uncoerced that competition occurs not so much by force or its threat, as due to a need to offer more of what the choosing party “demands”. The idea of biological markets thus sheds light on certain selective mechanisms, namely market effects in which *“members of one class can “force” members of another class to evolve traits that would have a negative effect on fitness in the absence of the cooperative interaction”* (Noë and Hammerstein, 1994, p. 2).

Along with other ecological forces, part of the evolutionary rationale of active cultural transmission might be a result of biological markets. Models and hypotheses akin to biological markets have already found applications in other arenas of evolutionary psychology, including the psychology of cooperation and mutualism (Frank, 1988; Baumard, 2010; André and Baumard, 2011). To our knowledge, Henrich and Gil-White (2001) first proposed that cultural abilities and knowledge could enter into a market-like exchange of what they called “information goods” and “prestige”. Based on previous anthropological observations (Barkow et al., 1975), they formulated a theory of human hierarchies, in which dominance and prestige hierarchies both differ and mix in the context of human hierarchical strategies. In humans, hierarchical status is attainable not only by use of force (i.e: the “dominance” strategy) or power, but also by demonstration of expertise in certain cultural domains, an ability that when is socially acknowledged is usually referred as “prestige” (Cheng et al., 2013). Since status tends to be associated with reproductive success and survival of the individual (Marmot, 2004), and since the use of force by way of sheer dominance was probably selected against during the evolution of our species (Boehm, 1999), pursuing competence and prestige might have been an advantageous reproductive strategy of primary importance in the history of our species. In what follows, we incorporate explicitly the modeling of that ecological force into the study of the evolution of cultural transmission.

## 2. Model 1: Absence of teaching

To present our modeling, as well as to underscore the necessity of introducing a perspective focused on biological markets, we begin by considering a simple producer/scrounger scenario with frequency dependence based on previous attempts at capturing basic processes in the evolution of social learning such as the influential pioneering work of anthropologist Alan Rogers (Rogers, 1988; Boyd and Richerson, 2004, for a review, see Aoki and Feldman, 2014).

We first suppose a minimal case in which there is no active teaching. Agents in the population can follow one of several strategies, each of which has the same baseline fitness, *W*_0_, in addition to the frequency-dependent fitness based on characteristics of the strategy. The strategies reproduce in the next generation with probability proportional to the fitness: if at time *t* there are *n*_*I*_ individuals with strategy *I* of fitness *W*_*I*_, then the number of individuals with strategy *I* at time *t* + 1 will be 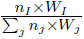. Such an idealization represents either the result of genetic evolution in an haploid panmictic asexual population or the dynamics resulting from social learning focused on the relative success of other strategies in the population.

In the simplest preliminary form of that scenario, a part of the population follows the strategy of individual learning. Those agents bear a cost *a* of learning individually. (This is a usual assumption in this type of models. Such a cost could represent either the cost of committing costly errors while learning by oneself, or the opportunity cost of investing time in individual learning, instead of something else.) At the same time, the strategy of individual learning also yields a benefit *α*. To simplify, we suppose here that agents who learn individually always discover an innovation of fitness value *α*. In the online appendix we show that if hiding the innovation is costly, then actively hiding individually acquired innovations is not an evolutionary stable strategy. Since this finding or discovery is partially observable, it is possible that other agents in the group will attempt to copy the solution by following a social learning strategy. We call agents who use that latter strategy “emulators”, and the process of social learning without evolved transmission of fitness enhancing behavior “inadvertent social transmission”.

To begin, we assume two conditions: First, only rarely does inadvertent social transmission produce perfect copies of behavior. Emulators who adopt the solution discovered by other agents thus benefit to the degree of *f* × *α* in which *f* < 1 is a transformation or “loss factor” associated with social learning (see Enquist et al., 2007 on the maladaptiveness of social learning). Second, we suppose that the easiness of social learning is directly proportional to the number of individual learners (Pagel, 2012). In our model, we codify that constraint by imposing a limited number of social learners *N*_*p*_ who can learn socially from a given individual learner. That condition is ecologically plausible, at least for a wide range of learning processes used to acquire certain techniques. Furthermore, it is easy to imagine that only a finite number of agents can have access to a given individual learner for the behavior to be adopted^1^.

The average fitness of an agent who learns socially is then dependent of the frequency of those who learn individually and that by the following rule:

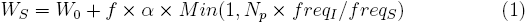

where *freq*_*X*_ is the proportion of strategy *X* in the population (*I* for individual learning, *S* for social learning), and *Min* denotes a selection of the minimal value between 1 and the effective proportion of emulators that can acquire the behavior given the number of individual learners in the population *N_p_* × *freq*_*I*_/*freq*_*S*_.

If *N*_*p*_ × *freq*_*I*_ > *freq*_*S*_, then all emulators can find a model to copy and their fitness will therefore equal *W*_*S*_ = *W*_*0*_ + *f* × *α*. In the opposite case, certain emulators can learn from a model but not all of them. The probability of learning socially is then *N*_*p*_ × *freq*_*I*_/*freq*_*S*_. If *f* × *α* < *α* −*a*, then the individual learning strategy is always more advantageous than the social learning one. At the same time, if *f* × *α* > *α* − *a*, then the number of social learners will tend to increase until *f* × *α* × *N*_*p*_ × *freq*_*I*_/*freq*_*S*_ = *α* − *a*, that is, to the point at which both strategies have the same fitness. At that equilibrium, it is the case that:

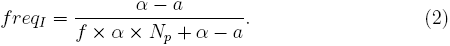

## 3. Model 2: Teaching in a biological market

### 3.1. Analytical model

A crucial feature of model 1 is that there is a maximum number *N*_*p*_ of emulators which can learn at a given time from one individual learner. At the equilibrium, not all emulators have the same kind of access to an individual learner. This is why there can be a market, the individual learners offering a privileged access to their skills (“selling”) in exchange for biological services. To introduce the possibility of teaching, we assume that agents who learn individually —with a frequency in the population *freq*_*I*_— can also follow a strategy by which they actively teach the acquisition of their technique. In addition to the cost of individual learning *a*, such a strategy will have a cost *t* linked to teaching. As with the previous model, we assume that there is a maximum *N*_*t*_ of individuals who can at once learn from a single teacher as “apprentices^2^”.

Another assumption of our model is that social learners who acquire the technique directly from the teacher will reproduce a perfectly efficacious copy of the teacher’s innovation. Although admittedly an idealization, the point is simply that, for this modality of technological learning, social learning without a teacher sometimes tends to produce a less fit solution than were there a teacher-apprenticeship relationship. Thus, if there is a teacher, then the fitness value of the socially learned technique becomes *α* instead of *f* × *α*.

However, individuals who learn socially from a teacher will recompense the teacher via deference and prestige mechanisms that have a cost *m* and that return *m* × *g* to the teacher. It seems reasonable to assume that most of the time *g* > 1, however our model does not strictly depend on that assumption.

For deference to evolve in social learning, its cost *m* must be less than the cost of individual learning *a*. Consequently, at its greatest value *m* is equal to *a*. For the evolution of teaching, the cost *t* of teaching thus has to be inferior to *N*_*t*_ × *g* × *a*.

Calculating the equilibrium state of the system is not straightforward, as both *freq*_*S*_ relative to *freq*_*I*_ and the value of *m* may evolve. Additionally, it could be that not all the individual have the same preference *m*.

One method is to look at the evolution of the frequencies and of *m* independently. We can start assuming that all the individuals in a population have a fixed preference *m*. We can write the fitness values for the teachers and apprentices and obtain their equilibrium frequencies for which their fitnesses are equal. Then, we assume that the frequencies are fixed, the whole population has still this preference *m*, except that there are mutants. If *N*_*t*_ *freq*_*I*_ < *freq*_*S*_, not all apprentices are matched with a teacher, and thus a mutant apprentice with a slightly higher *m* will be favored, and thus the preference *m* of the apprentices will evolve towards higher values, thus allowing the preference of the teachers *m* to also evolve towards higher values. Conversely, if *N*_*t*_ *freq*_*I*_ > *freq*_*S*_, not all teachers are paired with *N*_*t*_ apprentices, thus *m* is driven to decrease for the teachers, which then leads to a decrease in *m* for the apprentices. The next step is to study the effect of the change in *m* on the frequencies. For the initial *m*, the frequencies were such that apprentices and teachers had the same fitness. If *m* increases (respectively decreases), then teachers are less (respectively more) fit that apprentices, then the teacher’s frequency decreases (resp. increases). Thus the equilibrium point is:

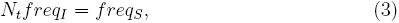

which is equivalent to:

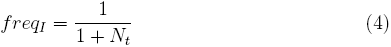

and:

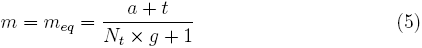

in which *m*_*eq*_ is the value of *m* at equilibrium.

By extension, another condition for the evolution of teaching is that *α > m_eq_*: an apprentice has to gain more through the acquired technique than the cost of deference. That condition is really constraining for teaching only at very high values of *a* or *t*. It is most reasonable that *g* is at least equal to 1, and *N*_*t*_ at least equal to 1. Thus for instance, if *a* and *t* remain less costly than *α*, then that condition is filled.

An interesting property of *m*_*eq*_ is that it is the *m* value maximizing the fitness of the population (see appendix). The evolutionary stable equilibrium is also the state of the system with the highest fitness. In our model, the so-called Rogers’ paradox (Aoki and Feldman, 2014) does not occur.

We considered here that all social learners have the same preference *m*, with *m* > 0, i.e. the social learners reward their teachers. But, there could be a distribution of preferences *m* in the population, and as in model 1, there could also be social learners (the “emulators”) who only try to copy without being taught, provided that *f* × *α* ≥ *α* − *m* (when *m* = *m*_*eq*_ (5), this condition is equivalent to 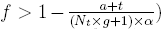. Interestingly, the presence of these emulators does not modify neither the equilibrium between the frequencies of the individual learning and apprentice strategies, nor the evolution of *m*. Even when *f* × *α* > *α* − *m*, apprentices are not driven to extinction by emulators. In other words, teaching may evolve even if social learning without teaching (“inadvertent social transmission”) is still an available and profitable strategy in the population. We can calculate the expected frequency of emulators: their frequency increases until there are not enough individual learners, so that *f* × *αN*_*p*_ *freq*_*I*_/*freq*_*emul*_ = *α* − *m*. This ultimately leads to:

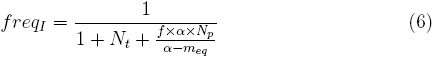

Even if deference is relatively costly, the apprentice strategy can be on par with the emulator strategy, because it enables a better access to individual learners who are a source of innovation. In fact, both strategies could still coexist with *f* =1 — albeit the larger *f*, the smaller the frequency of the apprentice strategy. As a result, the assumption that *f* < 1 is not necessary.

### 3.2. Simulation

The analytical calculations assumed mostly homogeneous *m* preferences, and evolution of the frequencies and *m* were considered separately. But both will actually evolve simultaneously, and if *m* mutates when the strategy is reproduced, preferences of teachers and apprentices cannot be exactly equal, because a mutation towards a slightly higher *m* for a teacher or a slightly lower *m* for an apprentice would lead to the inability to enter in a teaching relationship, thus decreasing the second-generation fitness.

To check that the system converges towards our analytical results, we coded an agent-based simulation. Each individual *j* is either an individual learner or a social learner, and attributes the reservation value *m*_*j*_ to teaching. Random pairs are formed between social learners who have not yet acquired the skill, and individual learners who have not yet taught to *N*_*t*_ social learners. If for a given pair, the reservation value *m* is smaller for the social learner, nothing occurs. But in the opposite case, the individual learner teaches the skill to the social learner at a price *m* which is taken as the average between the *m* values of the two individuals. (Any intermediate value between the two values would give similar results, see supplementary figure 12 in appendix.) This method of estimating the actual exchange value of *m* builds on the natural idea that there will be some form of bargaining between the two individuals. Pairs are formed until there is no possible additional interaction. Then the population is renewed, with new strategies taken at random proportionally to their fitness in the previous round, and the values *m*_*i*_ attributed to teaching in these strategies are copied with small random errors (to allow for the evolution of *m*). The frequencies of the different strategies and the average value of *m* tend to the state defined in equation (6) and (5), albeit with fluctuations around these values (see figure 2, and supplementary figures 2 to 13 in appendix). Having validated the results, we can now discuss them.

**Figure 1:**
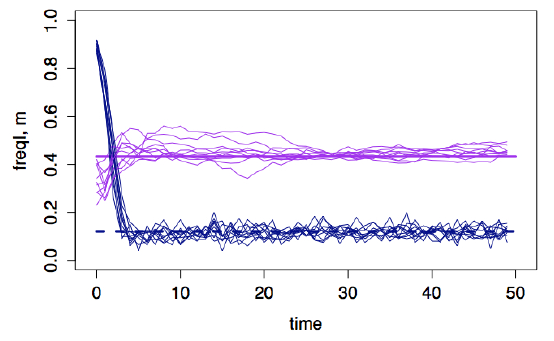
Simulation example. Frequency of individual learners (dark blue) and mean m values of interactions (light purple) for 10 different simulations, as a function of time (in generations). The horizontal thicker lines represent the predicted values of the frequency of individual learners (dark blue dashed line) and *m* (light purple solid line). For a population of 200 individuals, with *N*_*t*_ = 2, *N*_*p*_=5, *W*_0_=0.01, *α*=1, *a*=0.8, *f*=1, *g*=1, *t*=0.5, and *δm*=0.02. At the beginning of the simulations: 90% of the population are individual learners, a random *m* value is attributed to each individual, taken from a uniform distribution between 0 and 1.

**Figure 2:**
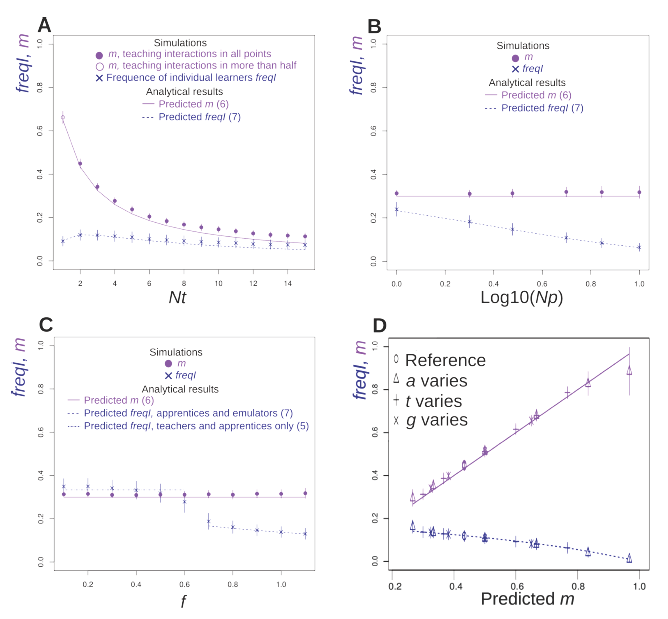
Dependence of the frequency of individual learners (dark blue) and average value of *m* in exchanges (light purple) with the different parameters. Results from simulations (symbols) taken averaged on 10 simulations, for generations 100 to 200 (see supplementary figure 1), and errors-bars represent the standard deviation. Theoretical curves: *m* (5) (solid purple lines), *freq_I_* with teachers, apprentices, and emulators (6) (dashed blue), with teachers and apprentices only (4) (dot-dashed blue), with individual learners and emulators only (2) (dotted blue). For all the simulations, the population is taken as 200 individuals, with the base fitness *W*_0_ = 0.01, the technique benefit *α* = 1, the typical mutational change on *m δ*_*m*_= 0.02, and when the interaction happens, *m* is taken as the average between the preferences of the two individuals (*coefshare* = 0.5). Initially 50% of the population are individual learners, with for all individuals, *m* taken at random between 0 and 1. Except if stated otherwise, the other parameters are *N*_*t*_ = 2, *N*_*p*_ = 3, *a* = 0.8, *t* = 0.5, *f* = 1, *g* = 1. Panel A: dependence on *N*_*t*_. Panel B: dependence on *N*_*p*_ (*t* = 0.1, *f* = 0.9). Panel C: dependence on *f* (*t* = 0.1). For panels A, B and C, *freq*_*I*_ in simulations is represented by triangles, and *m* in simulations is represented by a filled circle when there are exchanges in all the points used for computing the average values, and an empty circle when there are points for which there has not been any exchanges (but at least for half the points; *m* values for which there are exchanges in less than half the simulation points are not represented). Pannel D: as the dependence of *freq*_*I*_ on *t*, *g* and *a* is predicted to occur only through the value of *m* when there are teachers, apprentices and emulators, we represent the values of m and *freq*_*I*_ as a function of the predicted m for the points predicted to be within this regime. Supplementary figures 3, 4 and 5 show the dependence for each parameter individually.

### 3.3. Results

As confirmed by the numerical simulations, there are four different regimes, as summarized in table 1. Teaching is a stable strategy if the cost of teaching *t* is smaller than *N*_*t*_ × *g* × *a*. Teaching is clearly facilitated when there are more potential apprentices (*N*_*t*_) (Table 1 and supplementary figure 2), receiving deference provides a higher gain (*g*) (Table 1 and supplementary figures 5 and 9), and if learning the technique individually is costly (*a*) (Table 1 and supplementary figures 3 and 7). Interestingly, this condition does not depend on the characteristics of inadvertent social transmission (*N*_*p*_ and *f*) (Table 1 and panels B and C of figure 2). Profiting of “inadvertent social transmission” by emulators is a stable strategy if the loss in the technique value (1 − *f*)*α* is smaller than the cost of retributing a teacher (*m*_*eq*_) if there is teaching (Table 1 and figure 2C), or smaller than the cost of learning the technique individually (*a*) if there is no teaching (Table 1 and supplementary figure 7).

**Table 1:** Summary analytical regimes

|  |  |  |
| --- | --- | --- |
| $t <$<br>$N_t \times g \times a$ | $f > 1 - \frac{m_{eq}}{\alpha}$ | Teaching and emulating<br>(5) $m = m_{eq} = \frac{a+t}{N_t \times g + 1}$<br>(6) $freq_I = \frac{1}{1 + N_t + \frac{f \times \alpha \times N_p}{\alpha - m_{eq}}}$ |
| | $f < 1 - \frac{m_{eq}}{\alpha}$ | Teaching<br>(5) $m = m_{eq} = \frac{a+t}{N_t \times g + 1}$<br>(4) $freq_I = \frac{1}{1 + N_t}$ |
| $t >$<br>$N_t \times g \times a$ | $f > 1 - \frac{a}{\alpha}$ | Emulating<br>no $m$<br>(2) $freq_I = \frac{\alpha - a}{\alpha(1 + fN_p) - a}$ |
| | $f < 1 - \frac{a}{\alpha}$ | Individual learning only<br>No $m$<br>$freq_I = 1$ |

In the case of teaching, the value of the deference *m* at equilibrium increases with *a, t*, and decreases with *N*_*t*_ and *g*: deference has to be higher to offset a higher cost of individual learning and teaching, and the higher the number of apprentices per teacher and the higher the factor *g*, the less the deference cost per apprentice (table 1 and figures). The frequency of the teachers is such that there are *N*_*t*_ apprentices per teacher. If there are no emulators, then the teachers frequency depends only on *N*_*t*_. The teachers frequency is lower when there are emulators. It depends on *a*, *t*, *g* only through its dependence on *m* (table 1 and figure 2D). It decreases with *f* and *N*_*p*_: the more the emulators, the fewer the teachers and apprentices (table 1 and panels B and C of figure 2). It is maximum for some intermediate value of *N*_*t*_ (table 1 and figure 2A).

Another result is that, under a biological market of the type described here, for individual learning to be beneficial, it is sufficient that the cost of individual learning *a* is smaller than *α* × (*N*_*t*_*g* + 1) − *t*, which, except when *t* is large and *N*_*t*_ is small, is likely much larger than *α.* Hence, there are investments in skills for which benefits would not be sufficient in themselves, which thus become attractive because of the extra incentive linked to the sociability of teaching.

## 4. Model 3: Cumulative culture

Previous research has shown that social learning *per se* does not automatically lead to cumulative culture, that is, sustained evolution of ever increasingly adaptive cultural techniques (Enquist and Ghirlanda, 2007). In model 2, we have shown that the market for deference and prestige supports increased costs of innovations. Accordingly, we believe that taking those sorts of biological markets seriously can shed light on ecological forces active in the evolution of cumulative culture.

Until now, we have considered the skill to be fixed. Here, however, we consider a different model, in which the skill of value *α* can be improved by *δα* with probability *ϵ* when effort *r* is invested into innovation. We consider that at each time step, a new individual enters the population of size *N*, wheras the “oldest” individual dies. The entrance can represent either a birth — more realistically a child’s coming of age and being prepared for the apprenticeship — or a migration. If there is no active teaching, then the new individual copies the best skill in the population, of value *α*(*T*), but not perfectly. The individual will thus have a skill of value *f* × *α*(*T*), in which *T* represents the moment in time when the technique is copied by the newly arrived agent. We assume that individuals can recognize the best skill, and access the value of parameters *r*, *δα* and *ϵ*, as well as that innovation can occur only after the new individual entered the group and acquired its skill. This latter assumption represents the idea that some periods are more prone to innovation than others. The new individual can then decide whether to invest in innovation depending on whether *ϵ* × *δα* > *r*. If *ϵ* × *δα* < *r*, then no innovation is ever made, and, provided that *f* < 1, the skill will be completely lost in the population over time. If *ϵ* × *δα* > *r*, the skill will be improved upon by *δα* per each 1/*ϵ* new individuals on average. Population size matters (Kline and Boyd, 2010): if 1/*ϵ* > *N*, then the innovations will not occur often enough to preserve the skill in the population. If 1/*ϵ* < *N*, then the value of the skill over time will tend toward the point at which the imperfect copy and the innovation compensate: *α* = *δα*/(1 − *f*).

For active teaching, when the new individual enters the population, many potential teachers are available, meaning that there will be active teaching as long as *m* × *g* > *t*. Due to competition among teachers, *m* will tend to *t*/*g*. If *t*/*g* < (1 − *f*)*α*, then the new individual will prefer to learn the technique via active teaching instead of emulation, and thus end up learning the best skill *α* of the population. At that point, when choosing whether to invest in innovation, the individual will compare the investment cost *r* not only with the direct benefit (*ϵ* × *δα*), but also the direct benefit plus the benefit expected from teaching the innovation to the rest of the population. Since the new individual has a monopoly on the skill, other individuals in the population will recompense his or her teaching by a maximum of *m* = (1 − *f*)(*α* + *δα*), or even more if *N* > *N*_*p*_. Thus, the benefit expected from teaching is *Min*(*N*, *N*_*t*_)*g*(1 − *f*)(*α* + *δα*) − *t*.

In sum, populations with active teaching differ from those with only “inadvertent” social learning (emulation) in two ways. Because the skill can be learned more accurately, cumulative innovations are facilitated and the value of the skill can continue to increase. Furthermore, innovation is favored since its benefits might also derive from deference and prestige. Accordingly, in biological markets evolved teaching has the double effect of promoting cumulative culture, but also, and importantly, enhancing individual learning.

## 5. Discussion and conclusions

Modeling evolutionary social dynamics offers proof of the internal consistency of hypothesized evolutionary selective pressures (McElreath and Boyd, 2008). The models presented here thus corroborate the logical soundness of some intuitions previously formulated in purely verbal arguments (Henrich and Gil-White, 2001), as well as develop a well-articulated mathematical framework that adds to the paucity of models of the evolutionary milestone that is the evolution of teaching (Kline, 2015).

Reciprocity-based models are not usually well equipped to accommodate hierarchies and asymmetries as the ones that we describe in our model 2. Besides, reciprocity-based cooperation models usually focus more on the partner control aspect of repeated interactions than on partner choice, outside options, and active discrimination. We have shown that market effects can account for relevant dimensions of the sociability of teaching, such as the propensity to transmit fitness-enhancing information, as well as the evolution of deference and prestige. We believe that these important aspects of human social learning are better studied focusing on the supply and demand demographic dynamics of a biological trade, rather than on the standard reciprocity mechanism.

We have provided a partner-choice model of the evolution of teaching that focuses on the functional aspects of teacher-apprentice cooperation. This account cannot be per se an exhaustive evolutionary characterization of the emergence of teaching. Teaching is, after all, a complex ethological category that subsumes different —and presumably related— types of phenomena (Kline, 2015). Moreover, the models presented are not intended to be so much a realistic depiction of the actual evolutionary process, as an exploration of general ecological conditions for the evolution of teaching. However our work nevertheless points to possible evolutionary pathways, which perhaps because they had not been mathematically modeled, had not received much attention. One such possible paleoanthropological pathway is that the structure of communication and nonzerosumness inherent in the form of the basic apprenticeship system described here might have preceded —instead of followed— the evolutionary emergence of modern (i.e. Middle Paleolithic) human inventiveness (McBrearty and Brooks, 2000).

We have additionally shown that teaching can evolve under certain conditions. First, individual learning or learning without relying on others’ experience is costly. Second, certain techniques are constrained in terms of the number of individuals who can socially learn the technique from a single expert. Under those conditions, demographic dynamics could force social learners, who want to acquire the adaptive behavior discovered by individual learners, to pay a price in the form of deference. Furthermore, albeit unnecessary, the evolution of teaching is facilitated if, for learning certain techniques, social learning without explicit teaching —“eavesdropping” (Danchin et al., 2004)— yields an imperfect copying in which adaptive value can be lost. Crucially, genetic relatedness and parent-offspring nepotism (Castro and Toro, 2004) are not strictly necessary, either.

An important point that emerges from our work is that evolved teaching might be the mother of invention. In other terms, natural pedagogy and communication skills may precede, and not necessarily follow, the appearance of complex forms of culture. This dynamic runs counter to the perspective sometimes advanced holding that teaching evolved as a response to increasingly complex for novices, “opaque” cultural forms (Caldwell, 2015; Gergely and Csibra, 2006). According to that evolutionary hypothesis, complex cumulative culture necessarily preceded evolved teaching. However, as we show here, teaching might constitute an evolutionarily stable strategy even if the existing cultural forms are not opaque enough for novices: teaching could evolve when inadvertent social transmission (i.e. social learning without teaching) remains a thriving strategy in the population.

Undoubtedly, access to various forms of social learning cannot be controlled in a way to give rise to biological exchange markets: “eavesdropping” or inadvertent social facilitation could be the most frequent form of social learning in nature, perhaps even in humans. Nonetheless, in humans, important forms of technique acquisition can be reasonably controlled, even monopolized to some extent. For instance, ethnographic studies of stone-tool production (Stout et al., 2002) confirm that adult acquisition of certain sophisticated skills can be perceived as a form of transferable intellectual property. Such capacity for transmission is endowed with a form of authority often safeguarded and administered in a teacher-apprentice system via manifestations of personal commitment. In more modern settings, partner choice has widely been observed to be crucial to acquiring competence within organizations (Blau, 1964).

In contrast to nonhuman social learning, certain forms of human social learning are characterized by both the sophistication of cognitive mechanisms at work and the important constitutive role played by collaboration and nonzerosumness. These characteristics eventually give rise to apprenticeship structures (Waal, 2001; Sterelny, 2012). In this article, we have shown how those behavioral strategies can attain evolutionary equilibrium and persist in a population.

Naturally, not all forms of cultural diffusion rely on competence-based partner choice, a point that can hardly be overemphasized. However, some forms of human social learning depend far more on competence-based partner choice than others. Indeed, that aspect can help to explain the existence of several interesting regularities in the human psychology of competence assessment, admiration, and deference (Fessler, 2006).

At the proximate level, hierarchical tendencies of this sort are not entirely specific to humans. In fact, other animals have been observed to behave in ways consistent with the predictions of biological markets. In particular, nonhuman primates, such as chimpanzees (*Pan troglodytes*), have demonstrated an ability to discriminate possible partners based on their abilities (Melis et al., 2006). Experimental studies have furthermore shown how different species of primates can temporarily align their hierarchical behavior after individuals in their group have acquired some valuable cultural competence (see Stammbach (1988) for interesting work on Rhesus macaques). Grooming behavior has been shown to adapt to the supply and demand characteristics of a biological market in which at least one individual in a group has learned to use a tool to obtain a valuable shareable food (Fruteau et al., 2009). Indeed, it has even been suggested that, for some species, grooming could be a form of proto-currency in primate exchange markets (Barrett and Henzi, 2006).

In humans, considerable evidence points to the existence of both competence assessment and prestige-signaling behavior, the latter being a form of communicating that one excels in a given domain (Tracy and Matsumoto, 2008). Although the human ability to detect competence in given domains is certainly far from perfect (Mauboussin, 2012), it nevertheless works in a satisfying manner in many settings. Competence is assessed through both fast and slow processes of cognition. And quick it can be: in adults, judgments of competence can be made in as little as 100 milliseconds and those judgments are sometimes highly persistent and difficult to override (Fiske et al., 2007). Early on, children also begin to pay special attention to individuals judged as competent in a given domain (Keil et al., 2008). The current consensus maintains that children commonly use two different pathways to judge the reliability of an informant: one related to trust and benevolence, the other to competence and ability (Mascaro and Sperber, 2009; Harris, 2012).

Clearly, we have just scratched the surface of what biological trade models could offer for the modeling of social learning dynamics. It would be interesting to further explore the evolutionary dynamics linked to maladaptive biases in the human psychology of competence and prestige detection. For instance, we have not explored here the complex dynamics that could follow if social learners were to adopt the techniques and behaviors of other social learners who no longer track the environment through individual learning and innovation, yet might receive some form of social reward due to further transmitting a highly prized form of “knowledge”, even if it is directly ineffectual. Moreover, the amount of effort that cultural models put into teaching their apprentices, even if not genetically related to them, the diminishing fitness values of the technology if shared, or the reliability of the deference provided by the apprentices are all interesting features where genetic conflict and partner choice could be modeled fruitfully. We hope to encourage further work in this area.

Regarding important aspects of the evolution of cultural transmission, we have suggested that the partner-choice framework (Nesse, 2009) is better equipped than other theoretical frameworks that rely exclusively on either partner control or nepotistic genetic relatedness (Noë and Voelkl, 2013). The free-rider problem of fitness-enhancing cultural transmission, and cultural parental manipulation are largely by-passed by evolutionary systems such as those described in this article. Nearly a century ago, Lev Vigotsky characterized human social learning as an eminently cooperative activity. Biological market models can incorporate nonzerosumness of human social learning while at once accommodating important findings of the anthropology of deference and prestige, and revealing surprising evolutionary processes that lead to cumulative culture.

## Conflicts of interests

None

## Acknowledgements

Several. To be included later.

## Funding

During part of the preparation of this work, H.V. received support from a La Caixa Scholarship, as well as the “Estada breu d’un jove investigador convidat” program at the University of the Balearic Islands.

## Data availability

An online appendix including further analytic details, supplementary figures, and simulation code is provided.

## Footnotes

1 Mathematically, this condition helps to prevent singularities: without it, a single learning agent suffices in order for all social learners in a large population to be able to acquire the innovation (*N*_*pop*_ ≫ 1). However, the number of social learners would abruptly collapse (and become 0) when the proportion of individual learners decreases from 1/*N*_*pop*_ to 0. It is not incoherent to state that social learning is facilitated when the proportion of individual learners in the population is greater.

2 We make no assumption concerning the specific social configuration of the teacher-apprentice relationship, except that there is some nonzerosumness or collaboration in the basic terms described in the model.

